# Protective effect of plasma neutralization from prior SARS-CoV-2 Omicron infection against BA.5 subvariant symptomatic reinfection

**DOI:** 10.1101/2023.02.14.527605

**Authors:** Xiaosu Chen, Yanli Xu, Yan Xie, Weiliang Song, Ye Hu, Ayijiang Yisimayi, Fei Shao, Li Geng, Ying Wang, Hongmei Gao, Yansong Shi, Fei Gao, Ronghua Jin, Zhongyang Shen, Yunlong Cao

## Abstract

From December 2022 to January 2023, severe acute respiratory syndrome coronavirus 2 (SARS-CoV-2) infections caused by BA.5 and BF.7 subvariants of B.1.1.529 (Omicron) swept across mainland China. It is crucial to estimate the protective effect of the neutralizing antibodies generated by such mass infections against the next potential SARS-CoV-2 reinfection wave, especially if driven by CH.1.1 or XBB.1.5. Previously, we recruited and continuously followed a cohort of individuals that experienced Omicron BA.1, BA.2, and BA.5 breakthrough infections, as well as a control cohort with no history of SARS-CoV-2 infection. In the previously uninfected cohort, the total symptomatic infection rate surveyed during the outbreak was 91.6%, while the symptomatic reinfection rate was 32.9%, 10.5%, and 2.8% among individuals with prior Omicron BA.1, BA.2 and BA.5 infection, respectively, with median intervals between infections of 335, 225 and 94 days. Pseudovirus neutralization assays were performed in plasma samples collected from previously Omicron BA.1-infected individuals approximately 3 months before the outbreak. Results indicate a robust correlation between the plasma neutralizing antibody titers and the protective effect against symptomatic reinfection. The geometric mean of the 50% neutralizing titers (NT_50_) against D614G, BA.5, and BF.7 were 2.0, 2.5, and 2.3-fold higher in individuals without symptomatic reinfection than in those with symptomatic reinfection (*p* < 0.01). Low plasma neutralizing antibody titer (below the geometric mean of NT_50_) was associated with an enhanced cumulative risk of symptomatic reinfection, with a hazard ratio (HR) of 23.55 (95% CI: 9.23-60.06) against BF.7 subvariant. Importantly, neutralizing antibodies titers post one month after BF.7/BA.5 breakthrough infections against CH.1.1 and XBB.1.5 are similar to that against BF.7 from individuals with prior BA.1 infection while not experiencing a symptomatic BF.7/BA.5 reinfection (plasma collected 3 months before the outbreak), suggesting that the humoral immunity generated by the current BF.7/BA.5 breakthrough infection may provide protection against CH.1.1 and XBB.1.5 symptomatic reinfection wave for 4 months. Of note, the higher hACE2 binding of XBB.1.5 may reduce the protection period since the potential increase of infectivity.

## Main

From December 2022 to January 2023, severe acute respiratory syndrome coronavirus 2 (SARS-CoV-2) infections caused by BA.5 and BF.7 subvariants of B.1.1.529 (Omicron) swept across mainland China. The infection wave was rapid and widespread, impacting a large proportion of the population.^1,2^ It is crucial to estimate the protective effect of the neutralizing antibodies generated by such mass infections against the next SARS-CoV-2 reinfection wave, potentially driven by the newly-emerged Omicron variants CH.1.1, BQ.1.1, or XBB.1.5.

Previously, we recruited a cohort of individuals that experienced Omicron BA.1, BA.2, and BA.5 breakthrough infections in Jan-Feb, Mar-Jul, and Aug-Sep 2022 in Beijing and Tianjin, China, confirmed by polymerase chain reaction (PCR) (appendix p 5 and 8).^3-5^ The cohort was continuously followed, and plasma was collected every 3 months, until the mass infection wave in Dec 2022. A control cohort with no history of SARS-CoV-2 infection and similar vaccination profiles was also recruited in the same region (appendix p 8). Frequent PCR testing and epidemiological investigations based on the zero-COVID interventions ensured no additional infections prior to the current outbreak.

BF.7 was the predominant subvariant in Beijing and Tianjin during the December outbreak, with BF.7/BA.5 accounting for 68%/32% and 58%/42%, respectively.^6,7^ We found that in the previously uninfected cohort, the total symptomatic infection rate surveyed during December 2022 to January 2023 is 91.6%. Importantly, the symptomatic infection rates were 32.9%, 10.5%, and 2.8% among individuals with prior Omicron BA.1 (n = 231), BA.2 (n = 191) and BA.5 (n = 470) breakthrough infection, respectively, with median intervals between infections of 335, 225 and 94 days (Fig. 1A). Note that the confirmation of infection was based on survey, not by strict screening using PCR or antigen testing, which is a limitation of the study. Compared to previously uninfected individuals, prior Omicron infection was associated with reduced susceptibility to the current BF.7/BA.5 subvariant epidemic, with a relative protection rate of 64.1% (95% CI: 52.4%-73.1%), 88.6% (95% CI: 81.7%-93.1%) and 97.0% (95% CI: 94.3%-98.5%) for previous BA.1, BA.2 and BA.5 infections.

**Figure 1.**
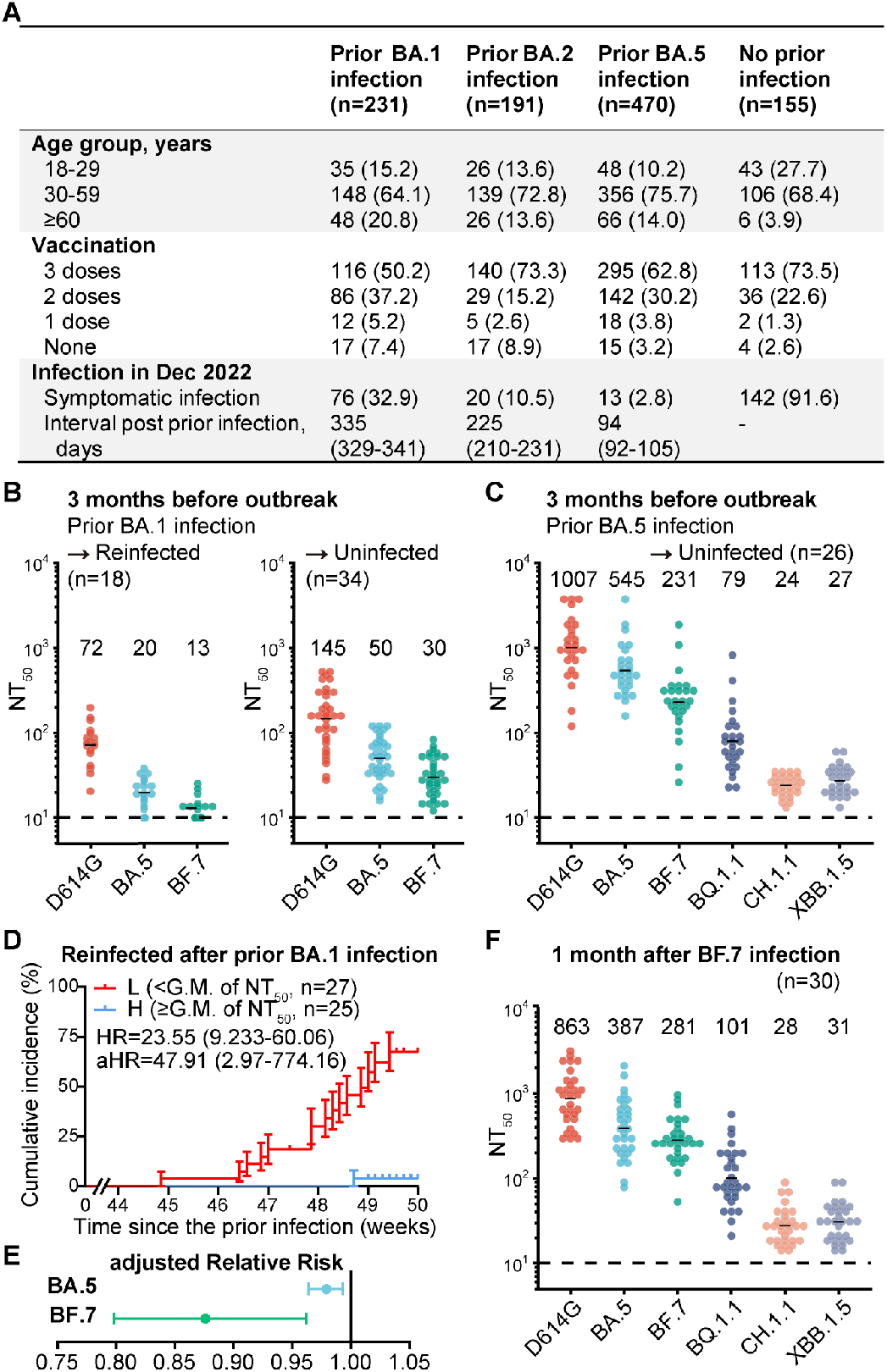
Protection by humoral immune responses in individuals with prior Omicron infection. (A) Demographic characteristics of individuals previously infected with Omicron BA.1/BA.2/BA.5 and those not infected with SARS-CoV-2, as well as the infection status in December 2022. Continuous variables were shown in median (interquartile ranges) and categorical variables were summarized as counts (percentages). (B) Plasma 50% neutralization titers (NT_50_) against SARS-CoV-2 D614G, BA.5, and BF.7 pseudovirus in those infected with prior BA.1 and reinfection (n = 18) and infected with prior BA.1 without reinfection (n = 34). The geometric mean titers are labeled, and dashed lines show the detection limits (NT_50_=10). (C) Plasma 50% neutralization titers (NT_50_) against SARS-CoV-2 D614G, BA.5, BF.7, BQ.1.1, CH.1.1, and XBB.1.5 pseudovirus in those infected with prior BA.5 without reinfection (n = 26). Plasma samples were collected approximately 3 months before the pandemic (1 month after prior infection). The geometric mean titers are labeled, and dashed lines show the detection limits. (D) Cumulative incidence of reinfection in previously BA.1-infected individuals (n = 52), stratified on plasma NT_50_ against BF.7 subvariant above or below the geometric mean titers. HRs (95% Confidence Intervals) from Cox regression model and Log-rank test. (E) Multivariate analysis of the efficacy of neutralizing antibody titers against symptomatic reinfection in previously Omicron-infected individuals by modified Poisson regression model. (F) Plasma 50% neutralization titers (NT_50_) against SARS-CoV-2 D614G, BA.5, BF.7, BQ.1.1, CH.1.1, and XBB.1.5 pseudovirus in those infected with BF.7 (n = 30). Plasma samples were collected approximately 1 month after prior infection. The geometric mean titers are labeled, and dashed lines show the detection limits (NT_50_=10).

We collected plasma samples from previously Omicron BA.1-infected individuals approximately 3 months prior to the current outbreak (appendix p 9). Pseudovirus neutralization assays were performed to estimate the correlation between neutralizing antibody titers and the protective effect against symptomatic reinfection. Importantly, the geometric mean of the 50% neutralizing titers (NT_50_) against D614G, BA.5, and BF.7 were 2.0, 2.5, and 2.3-fold higher in individuals without symptomatic reinfection than in those with symptomatic reinfection (*p* < 0.01, Fig. 1B). We also collected plasma samples approximately 3 months prior to the current outbreak from those previously infected with BA.5 subvariant. Plasma NT_50_ showed high levels of neutralization against Omicron BA.5 subvariants, which accounts for the rare symptomatic reinfection cases (Fig. 1C and appendix p 6). Low plasma neutralizing antibody titer (below the geometric mean of NT_50_) against BF.7 subvariant of the BA.1-infected cohort was associated with an enhanced cumulative risk of symptomatic reinfection, with a hazard ratio (HR) of 23.55 (95% CI: 9.23-60.06) (Fig. 1D). The same applies to NT_50_ against BA.5 subvariant (appendix p 7). Among those previously infected with Omicron, high neutralizing titers against BF.7 and BA.5 were indicative of a reduced probability of being symptomatically reinfected with BA.5 subvariants, with a relative risk (RR) of 0.876 (95% CI: 0.798-0.962) for BF.7 and a RR of 0.979 (95% CI: 0.964-0.993) for BA.5 (Fig. 1E and appendix p 10). Together, these results showed a robust correlation between the plasma neutralizing titers and the protective effect against Omicron symptomatic reinfection. Notably, the reason such a significant correlation was observed here could be owing to the extensive viral exposure of the population during the outbreak, which minimizes the chance that uninfected individuals were due to the lack of viral exposure.

To estimate the protective effect of the humoral immunity generated by the BA.5/BF.7 outbreak against the next potential SARS-CoV-2 reinfection wave, we also measured the neutralizing titers against BQ.1.1, CH.1.1, and XBB.1.5 in plasma collected from individuals one month after BF.7 or BA.5 breakthrough infections (Fig. 1C and F). The NT_50_ values of plasma from BF.7/BA.5 breakthrough infections against CH.1.1 and XBB.1.5 are similar to that against BF.7 of plasma from individuals with prior BA.1 infection and without experiencing a symptomatic BF.7/BA.5 reinfection during the December outbreak, and substantially higher than NT_50_ against BF.7 of plasma from those reinfected by BF.7/BA.5 subvariants.

Given that the plasma of individuals who avoided reinfection was collected 3 months before the outbreak and plasma from BF.7/BA.5 breakthrough patients were collected 1 month after infection, this suggests that the humoral immunity generated by the current BF.7/BA.5 breakthrough infection may provide at least 4 months of protection against the CH.1.1 and XBB.1.5 symptomatic reinfection wave, while the protection against a BQ.1.1 may last even longer. Of note, the higher hACE2 binding of XBB.1.5 may reduce the period of protection due to the potential increase of infectivity.^8^ Also, the lack of measurement of the mucosal and cellular immunity would add uncertainty to the estimation, which is a limitation of the study.

Y.C. is a co-founder of Singlomics Biopharmaceuticals and inventor of provisional patents associated with SARS-CoV-2 neutralizing antibodies. All other authors declare no competing interests.

## Supplementary Appendix

## Acknowledgments

We thank all the volunteers involved in this study for providing critical information and peripheral blood samples. This project is financially supported by the Ministry of Science and Technology of China and Changping Laboratory (2021A0201; 2021D0102), and the National Natural Science Foundation of China (32222030).

## Author contributions

Y.C. designed the study. X.C. and Y.C. wrote the manuscript. W.S., A.Y., F.S., and Y.C. performed the pseudovirus neutralization assays and analyzed the neutralization data. X.C., Y.Xu. and Y.Xie. performed the statistical analyses. Y.Xu., Y.Xie., L.G., H.G., S.Z., R.J., and Z.S. recruited and performed the follow-up in prior Omicron-infected cohorts and individuals with no SARS-CoV-2 infection history. X.C., Y.H., Y.W., and Y.S. collected plasma samples from previously Omicron-infected individuals.

## Methods

### Study design

This cohort study is aimed to investigate the protective efficacy of prior SARS-CoV-2 Omicron infection against reinfection of the BA.5* sublineage. Reinfection was defined as positive PCR tests or the development of clinical symptoms, occurring at least 90 days after confirmation of SARS-CoV-2 infection.^1^ We recruited previously infected patients from Tianjin First Central Hospital and Beijing Ditan Hospital. The following data were collected: the onset date, symptoms, laboratory tests, hospitalization of initial infection in the electronic medical record, COVID-19 vaccination history in the immunization planning information management system, and associated demographic information. The patients were infected with Omicron BA.1 (n = 316), BA.2 (n = 279) and BA.5 (n = 626) in January-February, March-July and August-September 2022, respectively. The infected subvariants of each cohort were confirmed either by sequencing or epidemiologically linked to other confirmed patients during regional outbreaks. A control group of 155 residents aged 18 years or older in Beijing or Tianjin, who had no history of SARS-CoV-2 infection until December 2022, was also recruited. They were verified by frequent free PCR tests and epidemiological surveys during 2022, with all PCR testing documented at healthcare and testing facilities.

The primary study endpoint was the emergence of symptomatic infections in a pandemic caused by Omicron BA.5/BF.7 subvariants after the adjustment of epidemic prevention policies after December 2022, that is, reinfection more than 90 days from the initial infection in the previously infected group, or the emergence of Omicron infection in the control group. We included autonomously reported positive cases that were symptomatic or epidemiologically linked but did not undergo PCR/antigen testing, as the cities under study had almost reached the point of saturation of infection since December 2022. The follow-up was conducted from December 25, 2022, to January 9, 2023, by telephone questioning and questionnaires. The average follow-up time was 201 days (range 95-366 days), with 350 days (range 331-366 days) for the BA.1 cohort, 243 days (range 165-305 days) for the BA.2 cohort, and 110 days (range 95-129 days) for the BA.5 cohort. All subjects with incomplete follow-up or partial missing data were excluded, resulting in a final study size of 892 cases in the previously infected group (n = 231 for BA.1, n = 191 for BA.2 and n = 470 for BA.5) and 155 cases in the control group. The efficacy of prior infection against the ongoing pandemic was defined as the ratio of the reduction in susceptibility between individuals with prior infection and those without prior infection to the rate of infection in individuals without prior infection, similar to the efficacy of vaccines.^2^

We collected peripheral blood samples from 52 individuals previously infected with BA.1 at a median of 241 days (interquartile range: 236-244 days) after initial infection and from 28 individuals previously infected with BA.5 at a median of 37 days (interquartile range: 32-40 days) after initial infection, approximately 3 months before follow-up. Additionally, we collected peripheral blood samples from 30 individuals previously infected with BF.7 at a median of 40 days (interquartile range: 36-47 days) after initial infection at the Fourth Hospital of Inner Mongolia in November 2022. However, due to the proximity of the onset of infection with BF.7 to the end of the study, follow-up and reinfection analyses were not performed in this cohort. Subsequently, we isolated plasma and performed pseudovirus neutralization assays to analyze the correlation between neutralizing antibody titers and symptomatic reinfection.

### Plasma isolation

Peripheral blood samples were collected from individuals in the cohort. The samples were processed by diluting the whole blood with PBS containing 2% FBS at a ratio of 1:1 and then subjected to Ficoll (Cytiva, 17-1440-03) gradient centrifugation. Plasma was collected from the upper layer and stored at -20 °C or below and was heat-inactivated before the experiments.

### Pseudovirus neutralization assay

Spike-pseudotyped virus of SARS-CoV-2 ancestral strain and Omicron BA.5, BF.7, BQ.1.1, CH.1.1, and XBB.1.5 sublineages carrying convergent mutations were prepared based on a vesicular stomatitis virus (VSV) pseudovirus packaging system. D614G spike virus (GenBank: MN908947 + D614G), BA.5 spike (T19I, L24S, del25-27, del69-70, G142D, V213G, G339D, S371F, S373P, S375F, T376A, D405N, R408S, K417N, N440K, L452R, S477N, T478K, E484A, F486V, Q498R, N501Y, Y505H, D614G, H655Y, N679K, P681H, N764K, D796Y, Q954H, N969K), BF.7 spike (BA.5 + R346T), BQ.1.1 spike (BA.5 + R346T, K444T, N460K), XBB.1.5 spike (BA.2 + V83A, del144, H146Q, Q183E, V213E, G252V, G339H, R346T, L368I, V445P, G446S, N460K, F486P, F490S, R493Q), BA.2.75 (BA.2+K147E, W152R, F157L, I210V, G257S, D339H, G446S, N460K, and R493Q) and CH.1.1 (BA.2.75 + R346T, K444T, L452R, F486S) spike plasmid were constructed into the pcDNA3.1 vector, and the pcDNA3.1-spike protein plasmids and G*ΔG-VSV (Kerafast) with firefly luciferase in place of VSV-G were transfected into 293T cells (American Type Culture Collection [ATCC], CRL-3216). The supernatants were discarded after 6-8 hours and replaced with a complete cell culture medium. Cells were cultured for 1 day, after which the pseudovirus was harvested from the supernatant, filtered (0.45-μm 195 pore size, Millipore), dispensed, and stored at -80°C for further use. Prior to use, multiple variants were diluted to the same number of copies.

Plasma samples were serially diluted and incubated with pseudovirus in 96-well plates in 5% CO_2_ at 37°C for 1 hour. Huh-7 cells (Japanese Collection of Research Bioresources, 0403) were seeded onto the plates and cultured for 1 day under 5% CO_2_ at 37°C. After removing half of the supernatant, the luciferase substrate (Perkinelmer, 6066769) was added to the plates and incubated for 2 minutes in darkness, followed by measuring the luminescence values using PerkinElmer Ensight (PerkinElmer, HH3400). The IC_50_ was calculated by fitting a logistic regression model with four parameters. All experiments are independently reproduced at least twice.

### Statistical Analysis

Unless otherwise specified, continuous variables were shown in medians and interquartile ranges (IQR). The Mann-Whitney U test was used to analyze the differences between the two groups. Categorical variables were summarized as counts and percentages. The efficacy of the prior Omicron infection against the ongoing epidemic of BA.5/BF.7 was estimated by the formula 100×(1−IRR), where IRR is the calculated ratio of incidence in the prior Omicron infection group to that in the control group. The Bayesian beta-binomial model with a threshold of 30% and a fairly wide prior probability distribution (95% confidence interval: 0.005-0.964) was used in calculating the efficacy of prior Omicron infection in preventing the ongoing epidemic.

The Log-rank test and a COX regression model were used to compare the risk of symptomatic infection among individuals with plasma-neutralizing antibody titers to SARS-CoV-2 D614G and Omicron subvariants above the geometric mean with those below. The modified Poisson regression model was used to estimate the relative risk while adjusting for age, gender, vaccination status, previously infected subvariants, and intervals between initial infection and the end of follow-up.

Various additional analyses and sensitivity analyses (adjusting multiple factors and different assumed distributions) were applied to validate the reliability of the main analysis. No interactions were investigated. The analyses were performed using the SPSS (ver22.0) or R software (ver4.2.2). Two-sided *p* < 0.05 was used in tests of significance.

### Study Oversight

This study was approved by the ethics committee of Tianjin First Central Hospital (ethics committee archiving No. 2022N151KY), the ethics committee of Beijing Ditan Hospital Capital Medical University (ethics committee archiving No. LL-2020-042-02), and the ethics committee of the Fourth Hospital of Inner Mongolia (Ethics committee archiving No. 202220). In accordance with the Declaration of Helsinki, written informed consent was obtained from each participant for the collection of their clinical information and blood samples for the purposes of the study and the publication of data generated by the study. All the authors contributed to data collection and analysis, discussion, and interpretation of the results. All the authors have read and approved the final manuscript.

## Supplementary figures

**Figure S1.**
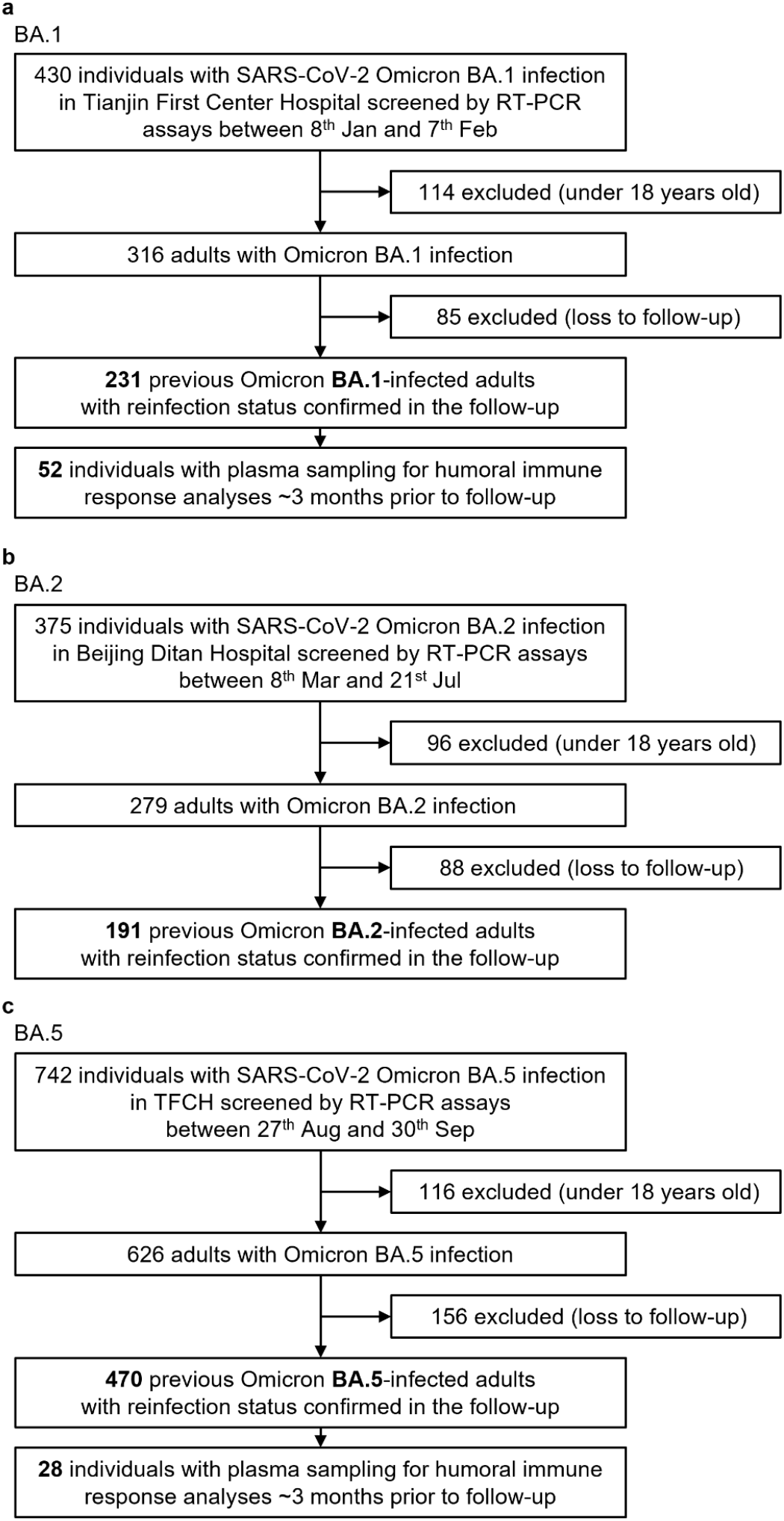
Flow chart of the recruitment and plasma sampling of individuals with a prior infection caused by Omicron BA.1 (a), BA.2 (b), and BA.5 (c).

**Figure S2.**
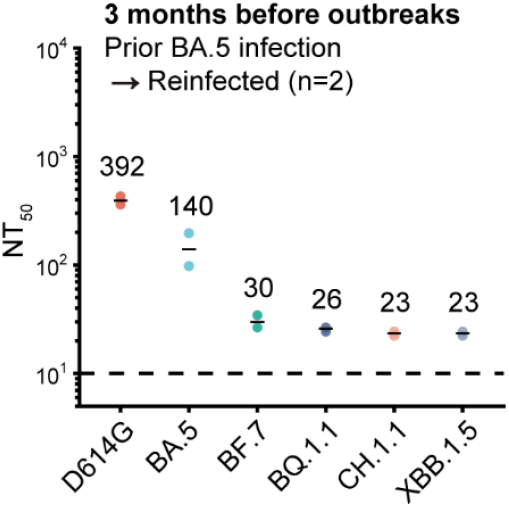
Humoral immune responses against D614G and subvariants of SARS-CoV-2 Omicron among previously BA.5-infected individuals with reinfection. Plasma 50% neutralization titers (NT_50_) against SARS-CoV-2 D614G, BA.5, BF.7, BQ.1.1, CH.1.1, and XBB.1.5 pseudovirus is displayed. The geometric mean titer is labeled, and dashed lines show the detection limits (NT_50_=10).

**Figure S3.**
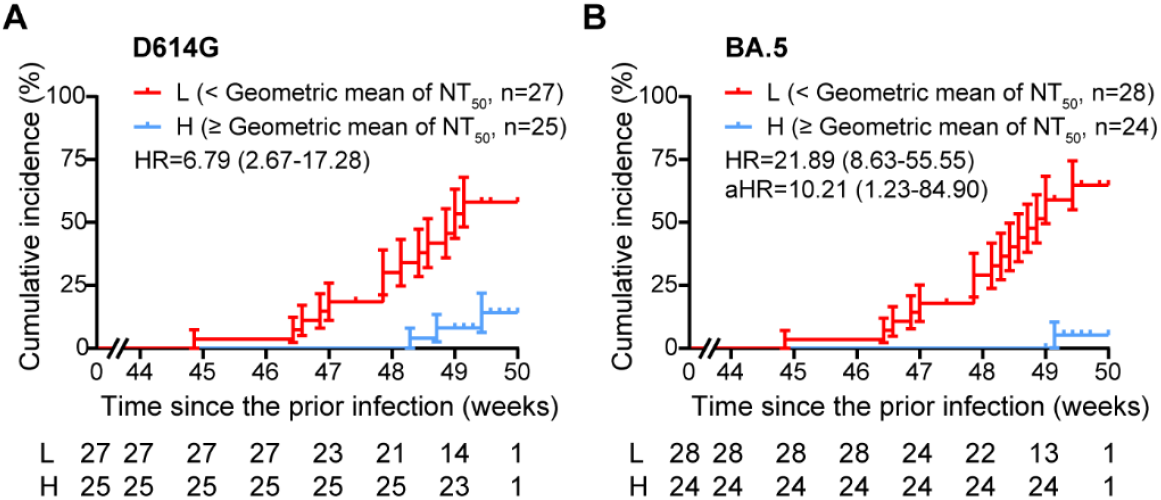
Cumulative incidence of reinfection in previously BA.1-infected individuals (n = 52), stratified on plasma NT_50_ against D614G (a), and BA.5 subvariant (b) above or below the geometric mean titers. HRs (95% Confidence Intervals) from Cox regression model and Log-rank test.

## Supplementary tables

**Table S1.**
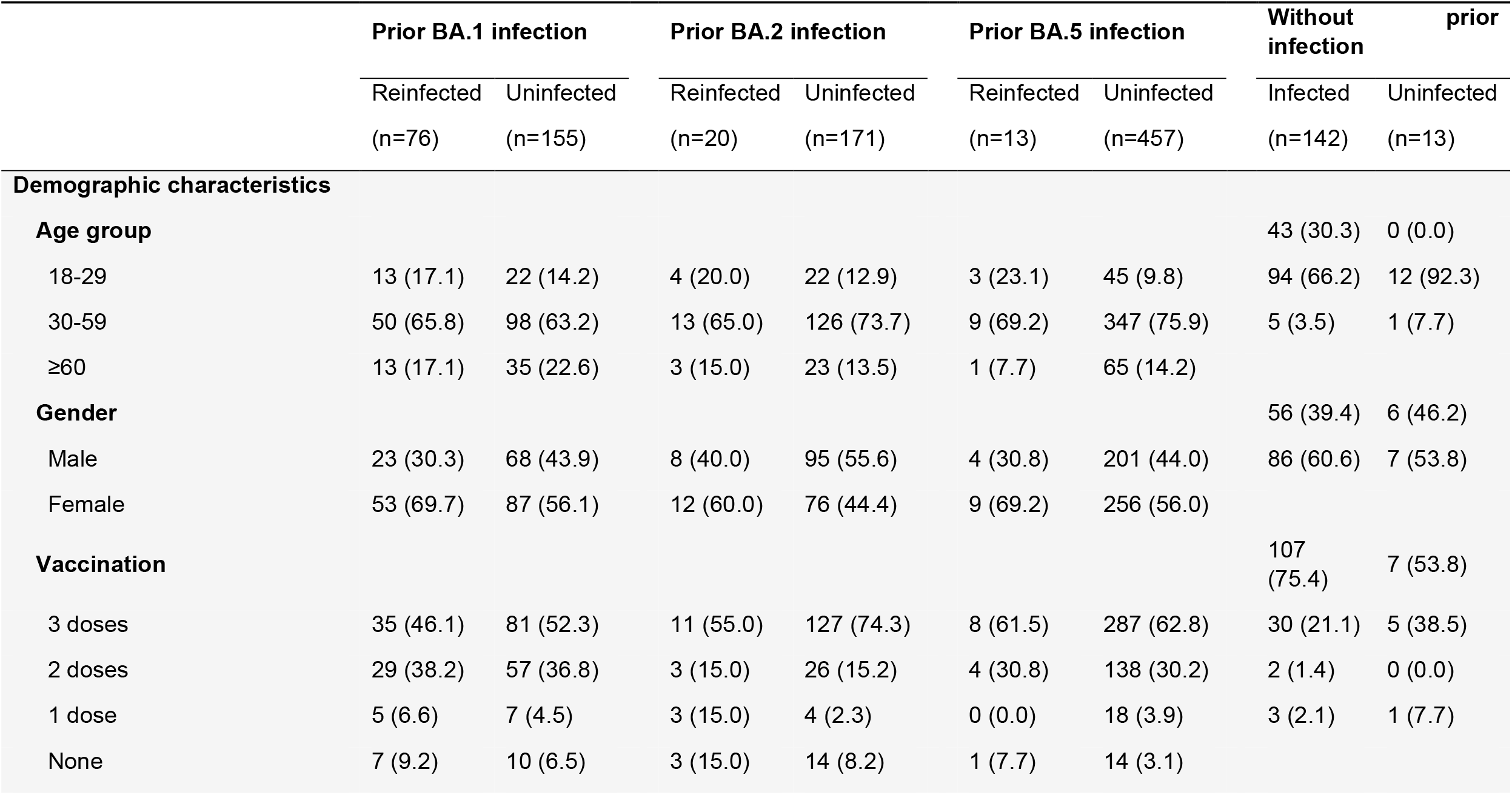
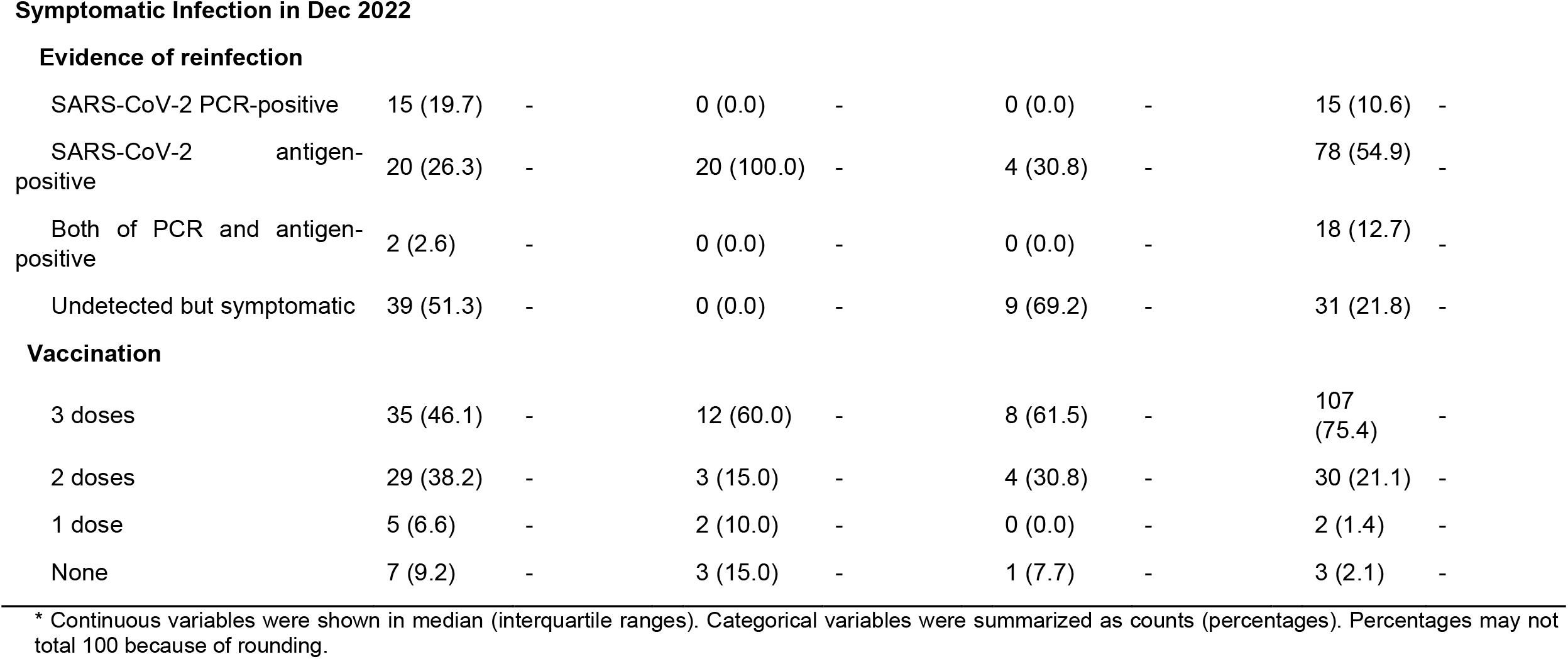
Characteristics of individuals with or without symptomatic infection in December 2022 in previously Omicron BA.1, BA.2, BA.5-infected cohorts or no prior SARS-CoV-2 infection cohort.

|  | Prior BA.1 infection |  | Prior BA.2 infection |  | Prior BA.5 infection |  | Without infection | prior |
| --- | --- | --- | --- | --- | --- | --- | --- | --- |
|  | Reinfected | Uninfected | Reinfected | Uninfected | Reinfected | Uninfected | Infected | Uninfected |
|  | (n=76) | (n=155) | (n=20) | (n=171) | (n=13) | (n=457) | (n=142) | (n=13) |
| **Demographic characteristics** |  |  |  |  |  |  |  |  |
| **Age group** |  |  |  |  |  |  | 43 (30.3) | 0 (0.0) |
| 18-29 | 13 (17.1) | 22 (14.2) | 4 (20.0) | 22 (12.9) | 3 (23.1) | 45 (9.8) | 94 (66.2) | 12 (92.3) |
| 30-59 | 50 (65.8) | 98 (63.2) | 13 (65.0) | 126 (73.7) | 9 (69.2) | 347 (75.9) | 5 (3.5) | 1 (7.7) |
| ≥60 | 13 (17.1) | 35 (22.6) | 3 (15.0) | 23 (13.5) | 1 (7.7) | 65 (14.2) |  |  |
| **Gender** |  |  |  |  |  |  | 56 (39.4) | 6 (46.2) |
| Male | 23 (30.3) | 68 (43.9) | 8 (40.0) | 95 (55.6) | 4 (30.8) | 201 (44.0) | 86 (60.6) | 7 (53.8) |
| Female | 53 (69.7) | 87 (56.1) | 12 (60.0) | 76 (44.4) | 9 (69.2) | 256 (56.0) |  |  |
| **Vaccination** |  |  |  |  |  |  | 107 (75.4) | 7 (53.8) |
| 3 doses | 35 (46.1) | 81 (52.3) | 11 (55.0) | 127 (74.3) | 8 (61.5) | 287 (62.8) | 30 (21.1) | 5 (38.5) |
| 2 doses | 29 (38.2) | 57 (36.8) | 3 (15.0) | 26 (15.2) | 4 (30.8) | 138 (30.2) | 2 (1.4) | 0 (0.0) |
| 1 dose | 5 (6.6) | 7 (4.5) | 3 (15.0) | 4 (2.3) | 0 (0.0) | 18 (3.9) | 3 (2.1) | 1 (7.7) |
| None | 7 (9.2) | 10 (6.5) | 3 (15.0) | 14 (8.2) | 1 (7.7) | 14 (3.1) |  |  |
| Evidence of reinfection |  |  |  |  |  |  |  |  |
| SARS-CoV-2 PCR-positive | 15 (19.7) | - | 0 (0.0) | - | 0 (0.0) | - | 15 (10.6) | - |
| SARS-CoV-2 antigen-positive | 20 (26.3) | - | 20 (100.0) | - | 4 (30.8) | - | 78 (54.9) | - |
| Both of PCR and antigen-positive | 2 (2.6) | - | 0 (0.0) | - | 0 (0.0) | - | 18 (12.7) | - |
| Undetected but symptomatic | 39 (51.3) | - | 0 (0.0) | - | 9 (69.2) | - | 31 (21.8) | - |
| Vaccination |  |  |  |  |  |  |  |  |
| 3 doses | 35 (46.1) | - | 12 (60.0) | - | 8 (61.5) | - | 107 (75.4) | - |
| 2 doses | 29 (38.2) | - | 3 (15.0) | - | 4 (30.8) | - | 30 (21.1) | - |
| 1 dose | 5 (6.6) | - | 2 (10.0) | - | 0 (0.0) | - | 2 (1.4) | - |
| None | 7 (9.2) | - | 3 (15.0) | - | 1 (7.7) | - | 3 (2.1) | - |

\* Continuous variables were shown in median (interquartile ranges). Categorical variables were summarized as counts (percentages). Percentages may not total 100 because of rounding.

**Table S2.** Characteristics of previously Omicron BA.1, BA.5, BF.7-infected individuals with plasma samples*.

|  | **Prior BA.1 infection cohort** |  | **Prior BA.5 infection cohort** |  | **BF.7 cohort** |
| --- | --- | --- | --- | --- | --- |
|  | Reinfected | Uninfected | Reinfected | Uninfected | (n=30) |
|  | (n=18) | (n=34) | (n=2) | (n=26) |  |
| **Demographic characteristics** |  |  |  |  |  |
| **Age, years** | 36 (33-42) | 46 (36-53) | 29 (26-31) | 37 (31-45) | 35 (30-41) |
| **Age group** |  |  |  |  |  |
| 18-29 | 3 (16.7) | 5 (14.7) | 1 (50.0) | 6 (23.1) | 7 (23.3) |
| 30-59 | 14 (77.8) | 23 (67.6) | 1 (50.0) | 19 (73.1) | 23 (76.7) |
| ≥60 | 1 (5.6) | 6 (17.6) | 0 (0.0) | 1 (3.8) | 0 (0.0) |
| **Gender** |  |  |  |  |  |
| Male | 7 (38.9) | 15 (44.1) | 0 (0.0) | 13 (50.0) | 9 (30.0) |
| Female | 11 (61.1) | 19 (55.9) | 2 (100.0) | 13 (50.0) | 21 (70.0) |
| **Vaccination** |  |  |  |  |  |
| 3 doses | 17 (94.4) | 31 (91.2) | 2 (100.0) | 24 (92.3) | 30 (100) |
| None | 1 (5.6) | 3 (8.8) | 0 (0.0) | 2 (7.7) | 0 (0.0) |
| **Reinfection in Dec 2022** |  |  |  |  |  |
| **Evidence of reinfection** |  |  |  |  |  |
| SARS-CoV-2 PCR-positive | 6 (33.3) | - | 0 (0.0) | - | - |
| SARS-CoV-2 antigen-positive | 6 (33.3) | - | 1 (50.0) | - | - |
| Undetected but symptomatic | 6 (33.3) | - | 1 (50.0) | - | - |
| **Interval between initial infection and follow-up, days** | 347 (344-348) | 346 (344-347) | 115 (115-116) | 116 (113-118) | - |
| **Interval between initial infection and sampling, days** | 241 (237-245) | 241 (236-244) | 24 (20-27) | 38 (32-40) | 40 (36-47) |
| **Interval between sampling and follow-up, days** | 103 (103-110) | 103 (103-110) | 92 (89-94) | 75 (75-86) | - |

\* Continuous variables were shown in median (interquartile ranges). Categorical variables were summarized as counts (percentages) Percentages may not total 100 because of rounding.

**Table S3.** Multivariate analysis of the efficacy of neutralizing antibody titers against symptomatic reinfection caused by BA.5/BF.7 subvariants in previously Omicron-infected individuals.

|  | Modified Poisson regression |  |  | Negative regression |  | Binomial |
| --- | --- | --- | --- | --- | --- | --- |
|  | RR (95% CI) |  | *p*-value | RR (95% CI) |  | *p*-value |
| **Age, year** | 0.993<br>(0.975-1.011) |  | 0.445 | 0.989<br>(0.965-1.013) |  | 0.355 |
| **Gender** |  |  |  |  |  |  |
| Female | 1.074<br>(0.606-1.903) |  | 0.807 | 1.149<br>(0.589-2.243) |  | 0.683 |
| Male | Reference |  |  | Reference |  |  |
| **Vaccination** |  |  |  |  |  |  |
| 3 doses | 1.158<br>(0.147-9.129) |  | 0.889 | 1.111<br>(0.146-8.428) |  | 0.919 |
| Unvaccinated | Reference |  |  | Reference |  |  |
| **Subvariant of prior infection** |  |  |  |  |  |  |
| BA.5 | 0.013<br>(0-26761379) |  | 0.691 | 0.002<br>(0-53042838) |  | 0.610 |
| BA.1 | Reference |  |  | Reference |  |  |
| **Intervals post initial infection, day** |  |  |  |  |  |  |
|  | 0.988<br>(0.905-1.079) |  | 0.789 | 0.979<br>(0.886-1.082) |  | 0.683 |
| **Neutralizing titers (NT<sub>50</sub>) against** |  |  |  |  |  |  |
| D614G | 0.999<br>(0.994-1.004) |  | 0.782 | 0.999<br>(0.994-1.004) |  | 0.697 |
| BA.5 | 0.979<br>(0.964-0.993) |  | 0.005 | 0.976<br>(0.960-0.991) |  | 0.002 |
| BF.7 | 0.876<br>(0.798-0.962) |  | 0.006 | 0.864<br>(0.793-0.941) |  | 0.001 |

